# Host gill attachment enables blood-feeding by the salmon louse (*Lepeophtheirus salmonis*) chalimus larvae and alters parasite development and transcriptome

**DOI:** 10.1101/815316

**Authors:** Erna Irene Heggland, Michael Dondrup, Frank Nilsen, Christiane Eichner

**Affiliations:** Department of Biological Sciences & Sea Lice Research Centre (SLRC), University of Bergen, Norway; Department of Informatics & Sea Lice Research Centre (SLRC), University of Bergen, Norway

**Keywords:** Salmon louse, ectoparasite, blood-feeding, hematophagy, gills, RNA-sequencing

## Abstract

Blood-feeding is a common strategy among parasitizing arthropods, including the ectoparasitic salmon louse (*Lepeophtheirus salmonis*), feeding off its salmon host’s skin and blood. Blood is rich in nutrients, among these iron and heme. These are essential molecules for the louse, yet their oxidative properties render them toxic to cells if not handled properly. Blood-feeding might therefore alter parasite gene expression. We infected Atlantic salmon with salmon louse copepodids and sampled the lice in two different experiments at day 10 and 18 post infestation. Parasite development and presence of host blood in their intestines were determined. We find that lice start feeding on blood when becoming mobile preadults if sitting on the fish body, however they may initiate in blood-feeding at the chalimus I stage if attached to gills. Lice attached to gills develop at a slower rate. Lice of similar instar age from gills versus lice from skin epidermis were analyzed for gene expression by RNA-sequencing in samples taken at day 10 for both experiments and at day 18 for one of the experiments. By differential expression analysis, we found 355 transcripts elevated in lice sampled from gills and 202 transcripts elevated in lice sampled from skin consistent in all experiments. Genes annotated with “peptidase activity” are among the ones elevated in lice sampled from gills, while in the other group genes annotated with “phosphorylation” and “phosphatase” is pervasive. Transcripts elevated in lice sampled from gills are often genes relatively highly expressed in the louse intestine compared with other tissues, while this was not the case for transcripts found elevated in lice sampled from skin. In both groups, more than half the transcripts are from genes higher expressed after attachment. In conclusion, blood-feeding results in an alteration in gene expression, and a premature onset of blood-feeding likely causes the parasite to develop at a slower pace.

## INTRODUCTION

The salmon louse, *Lepeophtheirus salmonis* (Krøyer, 1837) (Crustacea: Caligidae) and its Atlantic subspecies *L. salmonis salmonis* (Skern-Mauritzen et al., 2014), is an obligate ectoparasite of salmonid fish, such as the Atlantic salmon (*Salmo salar*). The parasite is of major concern for the aquaculture sector in the Northern Hemisphere, as it causes challenges for the industry with its high fecundity and resistance towards several chemotherapeutants (Aaen et al., 2015). The parasite life cycle consists of both planktonic and parasitic stages (Hamre et al., 2013; Johnson and Albright, 1991a). Upon hatching from a fertilized egg, the parasite is in the nauplius I stage. Thereafter, the salmon louse molts into the nauplius II stage, and further to the infective copepodid stage. Successive molting occurs on the host, first to the parasitic chalimus I and II. These stages are attached to the host by their elongated frontal filament (Bron et al., 1991; Gonzalez-Alanis et al., 2001), and are therefore immobile. Another molting renders the parasite mobile, as it is no longer secured by the frontal filament, but holds itself by using its cephalothorax as a suction cup. These stages are the preadult I and II and adult lice. Now, the parasite grazes on larger parts of its host, selecting its preferred feeding site and is causing greater damage to the fish (Bjørn and Finstad, 1998; Grimnes and Jakobsen, 1996). Progression of the salmon louse life cycle is temperature dependent, and at 10 °C, the time from fertilization to mature adult is approximately 40 (male) to 52 (female) days (Johnson and Albright, 1991b), or 38 (male) to 44 (female) days for the fastest developers (Hamre et al., 2019).

The alimentary canal of the salmon louse develops during the copepodid stage (Bron et al., 1993). The alimentary canal is composed of a mouthpart, an esophagus, a midgut, and a hindgut ending in a short rectum (Bron et al., 1993; Nylund et al., 1992). Vertebrate blood is a highly nutritious tissue fluid that is constantly renewed. Hematophagy (blood-feeding habit) is therefore a common strategy among parasitizing arthropods. The salmon louse diet is considered to consist of the skin and blood of its host (Brandal et al., 1976), and the blood-filled intestine is visible as a red line throughout the salmon louse body. Upon ingestion of blood, hematophagous parasites need to express genes encoding proteins that can manage the blood components. Blood is particularly enriched in proteins that contain the pro-oxidant molecules heme and iron. These are essential cofactors for the salmon louse, yet also highly toxic if not bound and detoxified by chaperones. Therefore, the alimentary canal needs to withstand, digest and absorb components of the food bolus. Trypsin-like enzymes (Johnson et al., 2002; Kvamme et al., 2004), a lipid transfer protein (Khan et al., 2017), a putative heme scavenger receptor (Heggland et al., 2019a) and the iron storage units of ferritin (Heggland et al., 2019b) are all expressed in the salmon louse midgut.

The distribution of copepodids on wild and farmed hosts shows that the preferred settlement site is on the fins and scaled body of the host (Bron et al., 1991). Some groups have reported the settlement of lice on gills as well, however this is considered rather uncommon (reviewed by Treasurer and Wadsworth (2004)). In laboratory trials, on the other hand, lice are often found on gills, although there still seems to be a higher preference for the fins and body (Bjørn and Finstad, 1998; Treasurer and Wadsworth, 2004). Copepodid gill settlement is therefore often considered an experimental artefact due to an altered host behavior during laboratory infestations (Treasurer and Wadsworth, 2004). Gill tissue in teleost fish is highly vascular, whereas skin epidermis is not. The chalimus frontal filament, appendages and mouth tube have been shown to not breach the basement membrane within the salmon skin (Jones et al., 1990), thus not reaching the dermal vascular layer. Salmon lice settling on gills might therefore be more prone to ingest a blood meal than those lice elsewhere on the host during early stages of attachment.

The genome of the Atlantic salmon louse is fully sequenced and high-throughput transcriptomics studies have been conducted under various experimental conditions using microarrays as well as sequencing. Examples of such experimental settings include host-parasite interactions on different hosts (Braden et al., 2017), hosts fed different diets (Sutherland et al., 2017), response to drugs (Sutherland et al., 2014), larval stress response (Sutherland et al., 2012), parasite sex differences (Poley et al., 2016), and development (Eichner et al., 2008). Recently, we have used RNA-sequencing (RNA-seq) to investigate patterns of gene expression during molting in parasitic larval stages of *L. salmonis* (Eichner et al., 2018). Transcriptome plasticity in response to hematophagy has been investigated in various arthropods for which controlled blood-feeding is possible. Arthropod species subjected to such controlled feeding trials include mosquitos (*Aedes* species (Bonizzoni et al., 2011; Bottino-Rojas et al., 2015; Huang et al., 2015), *Anopheles gambiae* (Marinotti et al., 2005)), the biting midge *Culicoides sonorensis* (Nayduch et al., 2014), and ticks (*Ixodes* species (Kotsyfakis et al., 2015; Perner et al., 2016). However, investigating transcriptional changes induced by a blood meal within the salmon louse is challenging, as no protocol for feeding lice *in vitro* exists. To overcome this limitation, equally developed lice of the same batch, infecting the same fish, were sampled from host body attachment sites with predicted differing access to blood.

In this study, we infected Atlantic salmon with salmon louse copepodids and sampled the lice on the 10th and 18th day post infestation (dpi), when the lice were in the chalimus I and chalimus II stage or had recently molted to the preadult I stage. Parasite settlement site and visible presence of host blood in louse intestines were determined. Transcriptomes of equally developed lice sampled from different locations (gills and the body/fins), representing lice with access to blood versus lice without access at 10 and 18 dpi, were examined by RNA-sequencing. Specific aims of this study were to investigate i) visible blood ingestion from various sampling locations, ii) development of lice from locations differing in blood access, and iii) differences in gene expression of immobile lice from locations with unequal access to blood.

## MATERIAL AND METHODS

### Animals

Atlantic salmon lice (*L. salmonis salmonis*) (Skern-Mauritzen et al., 2014) were raised on Atlantic salmon in tanks with seawater (salinity 34.5‰ and temperature 10 °C) (Hamre et al., 2009). A laboratory strain of *L. salmonis* called LsGulen (Hamre et al., 2009) was used. Fish were daily handfed commercial dry pellets and maintained according to Norwegian animal welfare regulations. Experiments conducted herein were approved by the governmental Norwegian Animal Research Authority (ID7704, no 2010/245410). Fish were anesthetized by a mixture of methomidate (5 mg/l) and benzocaine (60 mg/l) prior to handling. For sampling of early developmental stages of lice, fish were killed by a swift blow to the head. Salmon louse egg string pairs were incubated and hatched in incubators in a seawater flow through system (Hamre et al., 2009). Emerging copepodids were used to infect fish in 500-liter tanks. Copepodids between 4 and 14 days post hatching were used. A total of 34 or 37 fish in the two experiments respectively were infected with about 70 copepodids per fish. The amount of copepodids used was estimated as described by Hamre et al. (2009). Prior to infestation, the tank water was lowered and copepodids spread on the surface.

### Sampling of lice

At 10 and 18 dpi, fish were sacrificed and lice were removed with forceps and photographed. The gills were cut out and observed under a microscope. Any lice present were sampled, photographed and placed on RNAlater in individual tubes. Measurements of lice were done on photographs. Total length (TL) and cephalothorax length (CL) were measured as earlier described (Eichner et al., 2018, 2015) and determination of the developmental status as well as sex differentiation for chalimus II larvae was done using TL and CL measurements as described previously (Eichner et al., 2018, 2015). For RNA isolation prior to RNA-seq, lice were sorted into groups of equal developmental status within each group as described by Eichner et al. (2018), and between groups from different sampling locations (gills, skin). Five chalimus I or four chalimus II lice respectively were pooled together in one sample. Eight (Ex1 10 dpi), six (Ex2 10 dpi) or five (Ex2 18 dpi) replicates were sampled per group (lice sampled from gills, lice sampled from skin). RNA from both experiments sampled at 10 dpi and from one experiment sampled at 18 dpi was analyzed by RNA-seq.

### RNA isolation and sequencing

RNA was isolated as described before (Eichner et al., 2014). In brief, pools of four or five chalimus larvae were homogenized in TRI reagent and mixed with chloroform (both Merck). The water phase was withdrawn and further used with the RNeasy micro kit (Qiagen) for RNA isolation according to the manufacturer’s instructions. RNA was stored at −80 °C until being used. Library preparation and RNA-sequencing were conducted by the Norwegian Sequencing Centre, Oslo as previously described (Eichner et al., 2018). Briefly, sequencing libraries were prepared from 0.5 µg total RNA using the TruSeq stranded mRNA reagents (Illumina). Indexed libraries were blended into a single pool and sequenced during three runs of a NextSeq 500 instrument (Illumina) using 76 base pair (bp) single end reads. Image analysis and base calling were performed using Illumina’s RTA software version 2.4.11, and data were converted to fastq format using bcl2fastq version 2.17.1.14. Raw sequencing data has been deposited in the NCBI database under BioProject ID PRJNA577842.

### Data processing of RNA-seqencing data

Obtained sequences were quality controlled by FastQC v0.11.5 (Andrews, 2016). Reports were summarized using MultiQC 1.0 (Ewels et al., 2016). As reference genomes, we used a combination of the Ensembl Metazoa reference assembly of the nuclear genome (LSalAtl2s, http://metazoa.ensembl.org/Lepeophtheirus_salmonis) and the mitochondrial genome RefSeq sequence NC_007215.1 (Tjensvoll et al., 2005). The gene models from Ensembl Metazoa were further augmented with gene models derived from full-length sequences of LsFer1 and LsFer4 obtained by rapid amplification of cDNA ends (RACE) (Heggland et al., 2019b), by aligning the RACE consensus sequences against the nuclear assembly with GMAP (Wu and Watanabe, 2005). RNA-seq reads were aligned against the reference using the STAR aligner (Dobin et al., 2013). Then, alignments were sorted and indexed using SAM-tools (Li et al., 2009) and saved in BAM format. Technical replicates were merged prior to counting using the merge function in SAM-tools. RNA-seq reads and their overlap with annotated nuclear and mitochondrial transcripts were counted using the software featureCounts (Liao et al., 2014) with settings for strand-specific reverse stranded libraries.

Differential expression (DE) analysis was done with DESeq2 (Love et al., 2014) on raw counts using Galaxy (Giardine et al., 2005) under the Norwegian e-Infrastructure for Life Sciences (NeLS) platform (Tekle et al., 2018). Prior to DE analysis, all transcripts with less than four counts in all samples were removed. Venn diagrams were prepared using the BioVenn platform (http://www.biovenn.nl/) (Hulsen et al., 2008). Hierarchical clustering as well as GO annotation enrichment were performed in J-Express (Dysvik and Jonassen, 2001; Stavrum et al., 2008). GO terms were summarized in REVIGO (Supek et al., 2011).

### Transcript annotation

Protein-coding transcripts were annotated by running NCBI-Blast+, BlastP version 2.6.0+ (Altschul et al., 1990; Camacho et al., 2009) of their corresponding predicted Ensembl protein sequences against the GenBank (NR) (https://www.ncbi.nlm.nih.gov/) and SwissProt (Bairoch and Apweiler, 2000) databases. GO terms (full terms and GOslim annotation) and protein families (Pfam) were automatically assigned by InterProScan 5 (Jones et al., 2014).

## RESULTS

### Distribution and characteristics of lice

Upon termination all lice were removed from the salmon, and their settlement site and developmental stage and the visibility of an intestine filled with blood were assessed. Figure 1 depicts the distribution of different developmental stages and instar ages of lice on the host at 10 and 18 dpi. At 10 dpi, most lice are attached to the fins, but there is also a high proportion of lice on the body and gills. At 18 dpi, however, the highest proportion of lice is found on the body of the salmon. Different stages are distributed differently between sites.

**Figure 1.**
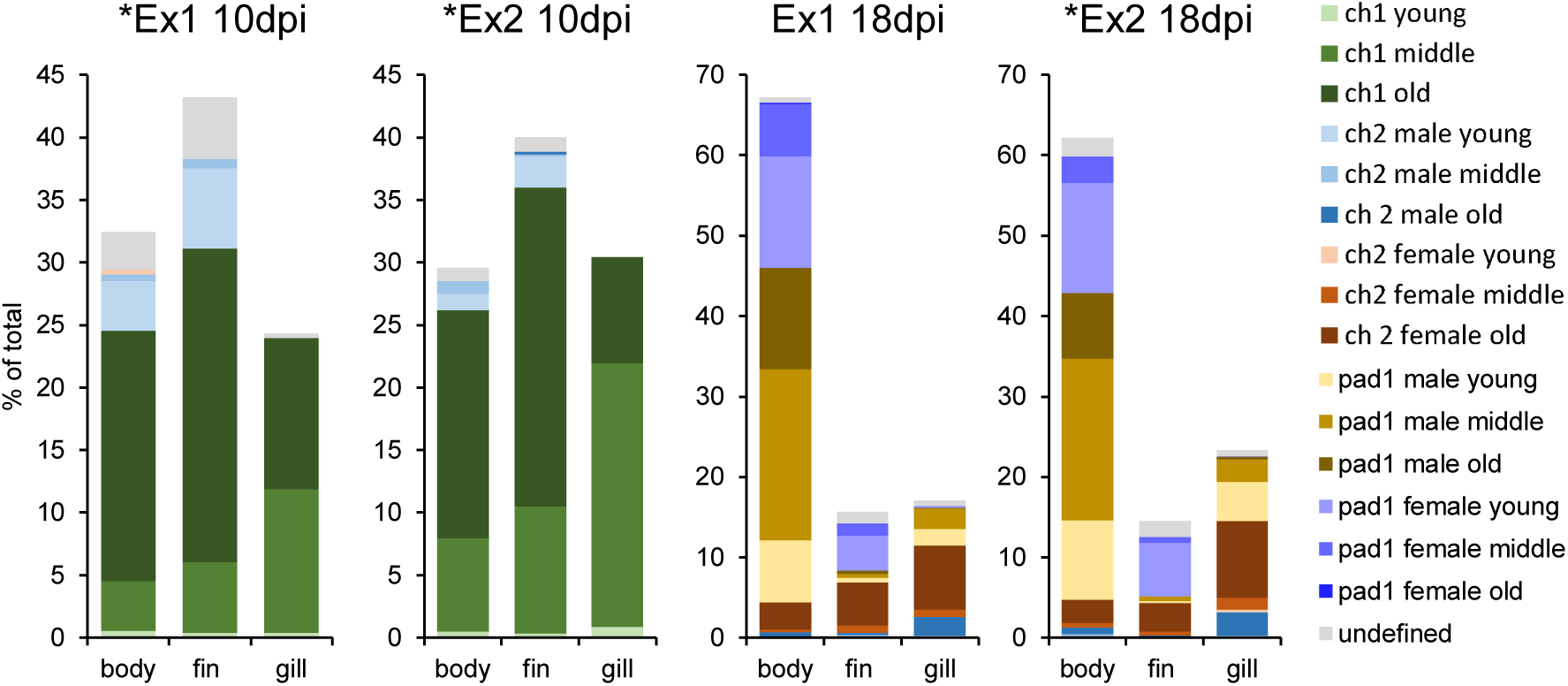
Distribution of different stages and instar ages of lice on fish at 10 days post infestation (dpi) and at 18 dpi sampled from the fish body, fins and gills in both experiments (Ex1 and Ex2). Lice instar ages were defined on photographs as described in the main text. Transcriptome sequencing was performed for lice from Ex1 10 dpi, Ex2 10 dpi (both chalimus I old), and Ex2 18 dpi (chalimus II old) marked with *.

At 10 dpi, the highest proportion of lice is in the chalimus I stage (83% and 95% in Ex1 and Ex2 respectively) while the remaining lice are chalimus II. The chalimus II larvae (mainly males) are found on the body and fins, but none on gills. Here, we rather find chalimus I (higher proportion of middle than old) larvae (Figure 1, Table 1).

**Table 1.**
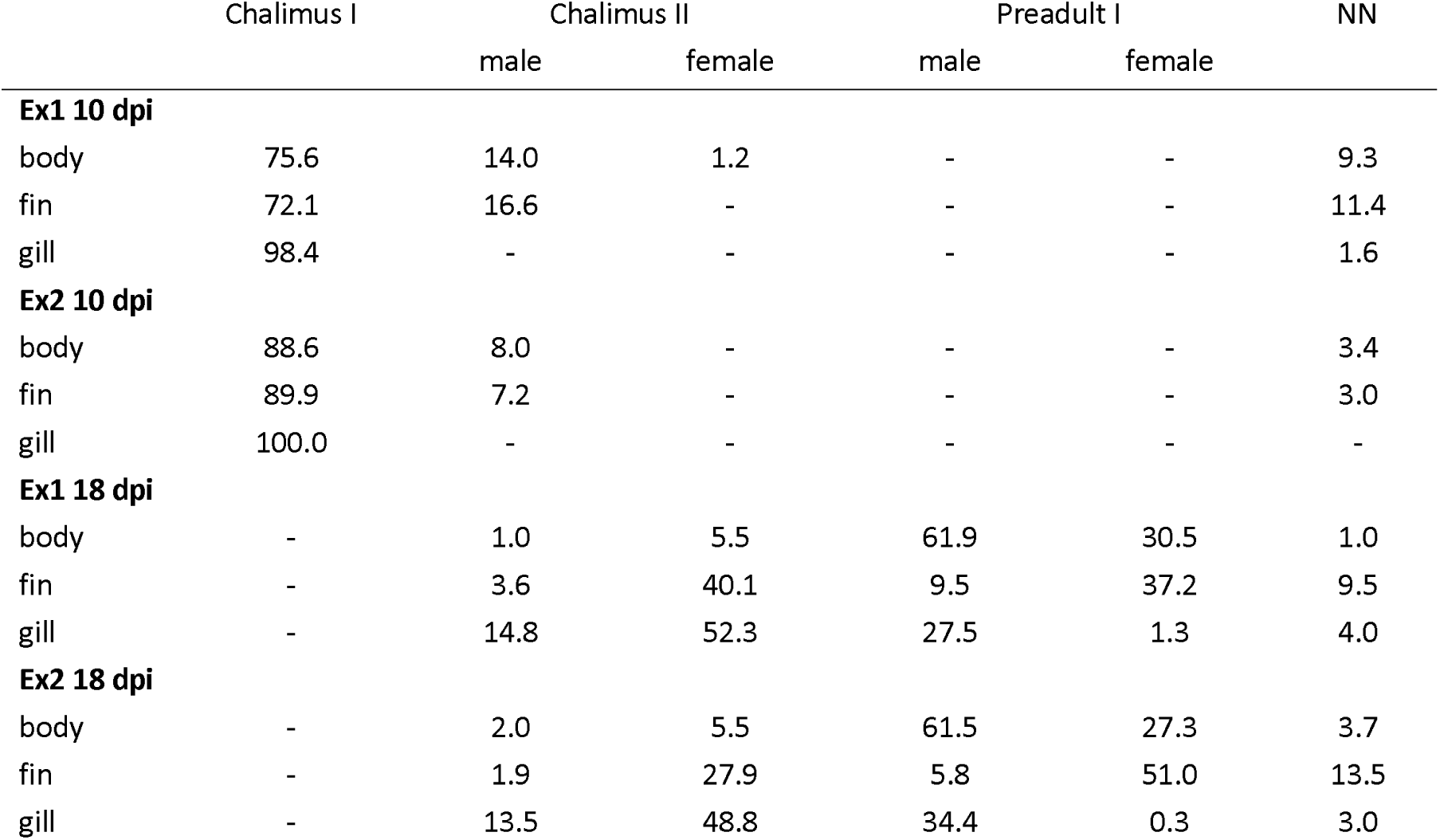
proportion of different stages and sexes (for Chalimus II and preadult I) from all fish sampled at 10 days post infestation (dpi) and 18 dpi in Experiment (Ex) 1 and 2. NN = undefined

|  | Chalimus I | Chalimus II |  | Preadult I |  | NN |
| --- | --- | --- | --- | --- | --- | --- |
|  |  | male | female | male | female |  |
| **Ex1 10 dpi** |  |  |  |  |  |  |
| body | 75.6 | 14.0 | 1.2 | - | - | 9.3 |
| fin | 72.1 | 16.6 | - | - | - | 11.4 |
| gill | 98.4 | - | - | - | - | 1.6 |
| **Ex2 10 dpi** |  |  |  |  |  |  |
| body | 88.6 | 8.0 | - | - | - | 3.4 |
| fin | 89.9 | 7.2 | - | - | - | 3.0 |
| gill | 100.0 | - | - | - | - | - |
| **Ex1 18 dpi** |  |  |  |  |  |  |
| body | - | 1.0 | 5.5 | 61.9 | 30.5 | 1.0 |
| fin | - | 3.6 | 40.1 | 9.5 | 37.2 | 9.5 |
| gill | - | 14.8 | 52.3 | 27.5 | 1.3 | 4.0 |
| **Ex2 18 dpi** |  |  |  |  |  |  |
| body | - | 2.0 | 5.5 | 61.5 | 27.3 | 3.7 |
| fin | - | 1.9 | 27.9 | 5.8 | 51.0 | 13.5 |
| gill | - | 13.5 | 48.8 | 34.4 | 0.3 | 3.0 |

At 18 dpi, we find mostly preadult I lice (75% or 74% in Ex1 and Ex2 respectively). On the body and fins, most lice were at the preadult I stage, while on the gills, most lice were in the chalimus II stage. Preadult I females are found on the body and fins, but nearly none on the gills (<1%). On gills, we rather find a higher proportion of old chalimus II females. There is a higher proportion of preadult I male lice found on the body compared with the fins. On the fins, most of the preadult I lice are young females. (Figure 1, Table 1)

For preadult I lice from 10 or 9 fish in Ex1 and Ex2 respectively also the presence of a frontal filament was investigated. Presence of a frontal filament and a visible blood filled intestine with respect to the settlement site are summarized in Table 2. The majority of the preadult I lice were located on the host body, and the minority was located on the gills. The preadult I lice on the gills, however, were more often secured by their frontal filament. Of the preadult I lice still attached by the filament, only the ones on the gills had a blood-filled intestine (Figure 2c). None of the lice on the fins had a blood-filled intestine, and on the host body, only the mobile lice had apparently fed on blood. Additionally, we found both chalimus I (Figure 2a) (at 10 dpi) and chalimus II (Figure 2b) larvae attached to the gills that had fed on blood, whereas lice of the same age on the fins and body had no visible blood in the intestine.

**Table 2.**
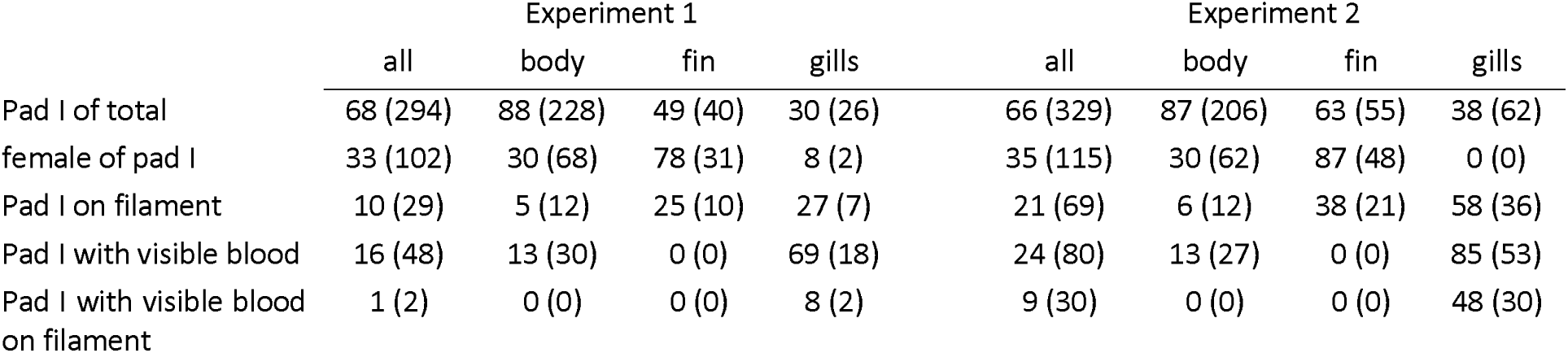
Preadult (pad) I lice were investigated in more detail of 10 and 9 fish in Experiment 1 and 2 respectively. The proportion and absolute numbers (in brackets) of preadult I lice on the different body parts, the proportion of females of these as well as the proportion of preadult I lice with visible blood in the intestine or with filament or with both are shown.

|  | Experiment 1 |  |  |  | Experiment 2 |  |  |  |
| --- | --- | --- | --- | --- | --- | --- | --- | --- |
|  | all | body | fin | gills | all | body | fin | gills |
| Pad I of total | 68 (294) | 88 (228) | 49 (40) | 30 (26) | 66 (329) | 87 (206) | 63 (55) | 38 (62) |
| female of pad I | 33 (102) | 30 (68) | 78 (31) | 8 (2) | 35 (115) | 30 (62) | 87 (48) | 0 (0) |
| Pad I on filament | 10 (29) | 5 (12) | 25 (10) | 27 (7) | 21 (69) | 6 (12) | 38 (21) | 58 (36) |
| Pad I with visible blood | 16 (48) | 13 (30) | 0 (0) | 69 (18) | 24 (80) | 13 (27) | 0 (0) | 85 (53) |
| Pad I with visible blood on filament | 1 (2) | 0 (0) | 0 (0) | 8 (2) | 9 (30) | 0 (0) | 0 (0) | 48 (30) |

**Figure 2.**
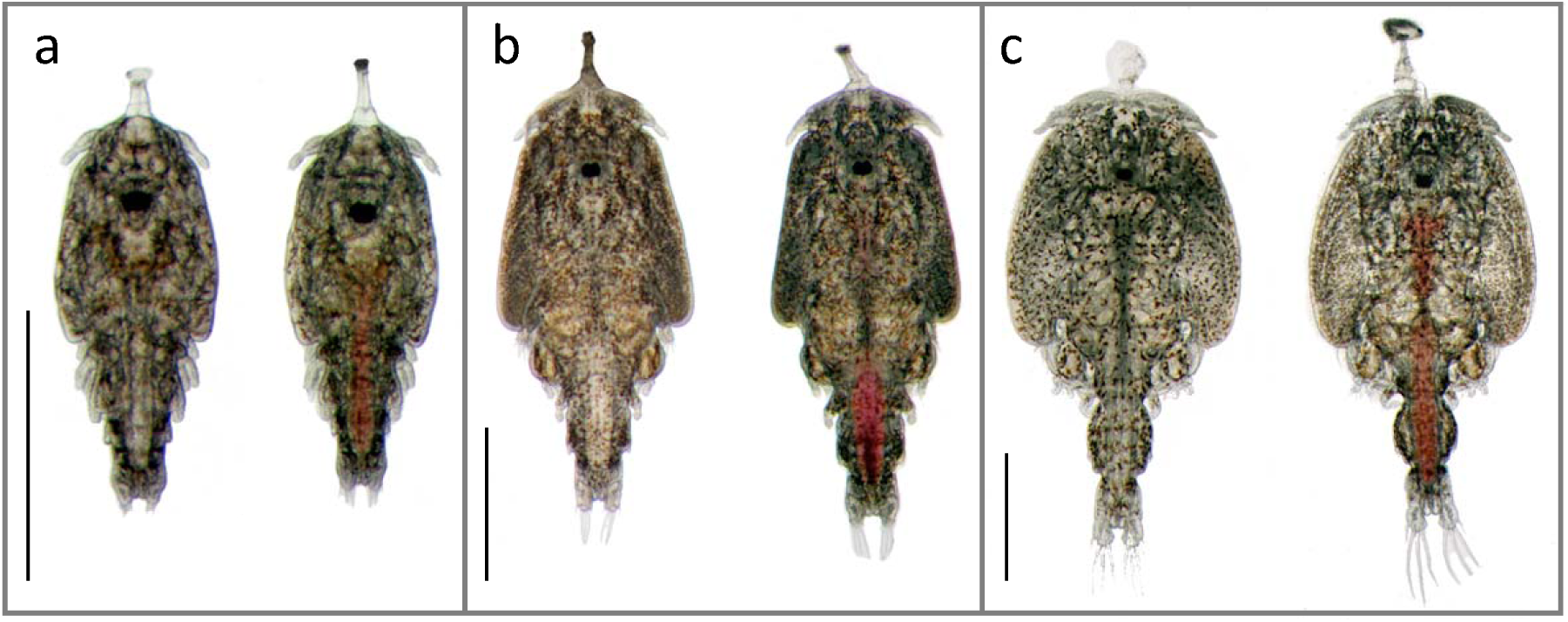
Photographs of salmon lice with (right) and without (left) a blood-filled intestine. Chalimus I larvae sampled 10 days post infestation (a). Chalimus II larvae sampled 18 days post infestation (b). Preadult I lice on frontal filament sampled 18 days post infestation (c). The lice with blood-filled intestine were sampled from the gills, and the others were sampled from the skin of their host. Scale bars = 1 mm.

### Effect of gill settlement on gene expression in chalimus larvae

In order to determine the effect of gill settlement on the gene expression in chalimus larvae, RNA-sequencing of pooled individuals of equal development was performed. All counts per million (CPM) values can be found in Supplementary Table S1. The overall gene expression of the individual samples in comparison with chalimus I and chalimus II larvae of different instar age (data taken from Eichner et al. 2018) is shown in a correspondence analysis (CA) plot (Figure 3c).

**Figure 3.**
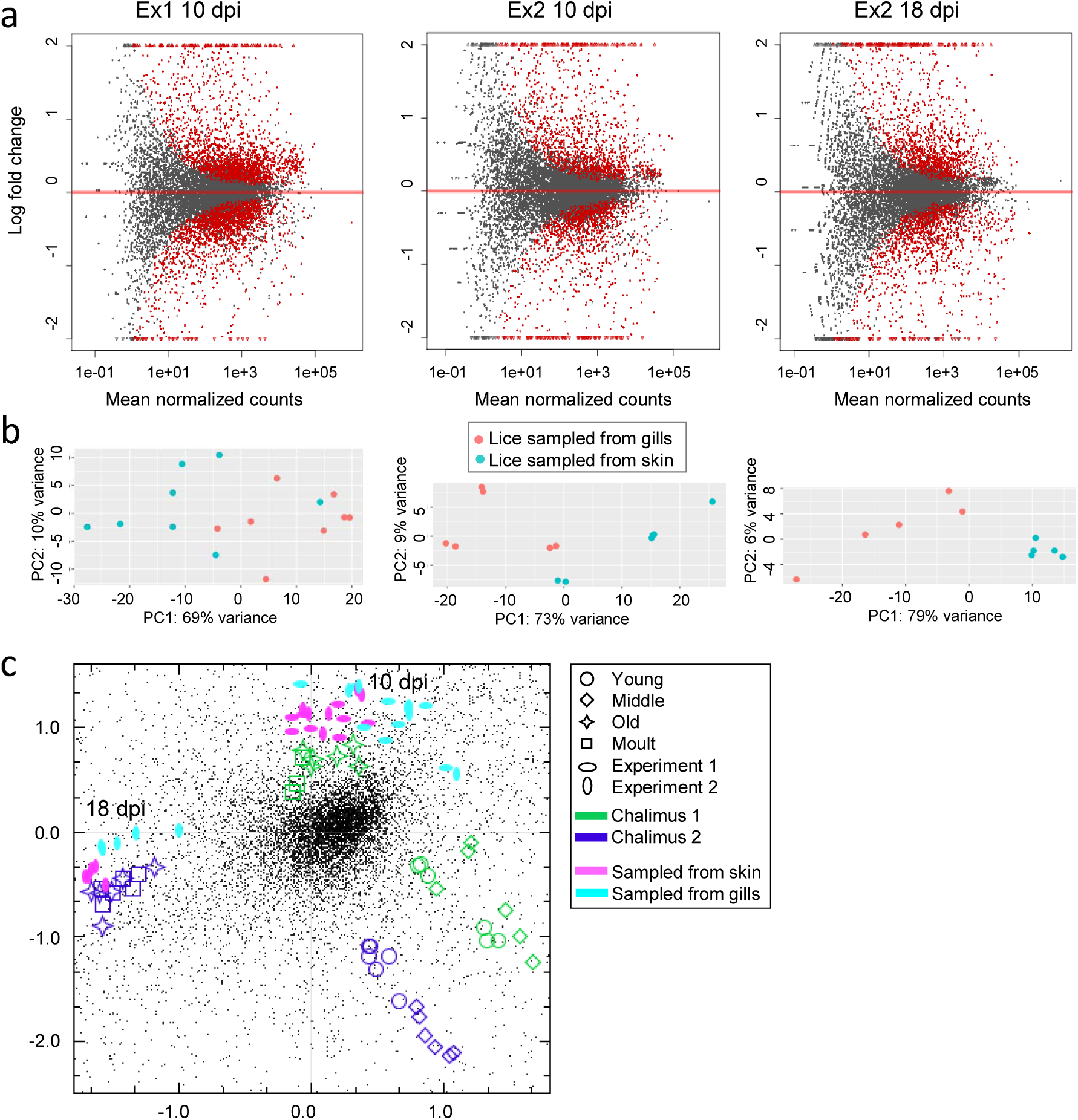
a) MA plot comparing the different conditions (lice sampled from gills versus lice sampled from skin at 10 days post infestation (dpi) and 18 dpi respectively) in the samplings: Experiment (Ex) 1, sampled at 10 dpi, Ex2 sampled at 10 dpi and Ex2 sampled at 18 dpi. The average log intensity for a dot in the plot (A) is shown on the x-axis and the binary logarithm of the intensity ratio (M) is shown on the y-axis. b) Principle component analysis (PCA) plots of the same data. c) Correspondence analysis (CA) plot showing the overall gene expression of the samples analyzed in this study (pink and turquoise dots) in comparison with other chalimus I and chalimus II larvae divided into various instar ages taken from Eichner et al. (2018).

All lice from this study sampled at 10 dpi are clustering together with chalimus I larvae sampled directly before molting as well as molting ones (old, molt) from Eichner et al. (2018) and all lice sampled at 18 dpi from this study are clustering with chalimus II lice sampled directly before molting (Eichner et al., 2018). Lice sampled from gills and lice sampled from skin differed also slightly in their overall gene expression. Lice from Ex1 at 10 dpi cluster together with lice from the respective group (from gills or from skin) of Ex2, showing lice sampled at 10 dpi in the different experiments are composed of comparable batches of lice. DE analyses were performed for each sampling separately. MA plots as well as a principle component analysis (PCA) for each sampling are shown in Figure 3a and b. A list of all genes with log2 fold changes and false discovery rate (FDR) adjusted p-values (padj) for each sampling can be found in Supplementary Table S2. 5878 genes were differentially expressed in at least one of the samplings (Supplementary Table S3).

Most DE genes were found in Experiment 1, 10 dpi (2188 or 2015 up-regulated in gill or skin samples respectively). In Experiment 2, 10 dpi only 1112 or 1081 transcripts respectively were found, of which 79% or 68% respectively are overlapping with the ones found in Ex1 10 dpi. DE genes found at 18 dpi are less overlapping with DE genes found in Ex1 at 10 dpi. Only 43% or 32% respectively of the genes found here are overlapping with genes from the respective groups in Ex1 10 dpi and 35% or 28% respectively are overlapping with Ex2 10 dpi (Figure 4). 616 genes are DE in all three samplings, of which 355 are elevated in lice samples from gills and 202 are elevated in lice samples from skin (Supplementary Table S6 and S7), while 59 are significant different, but regulation directions are differing between time points (31 elevated in lice from gills at 10 dpi but lower at 18 dpi, 24 the other way around and 4 are differing between lice sampled at day 10 of the two different experiments) (Supplementary Table S8). Transcripts solely regulated at either 10 dpi or 18 dpi are listed in Supplementary Table S4 and S5, respectively.

**Figure 4.**
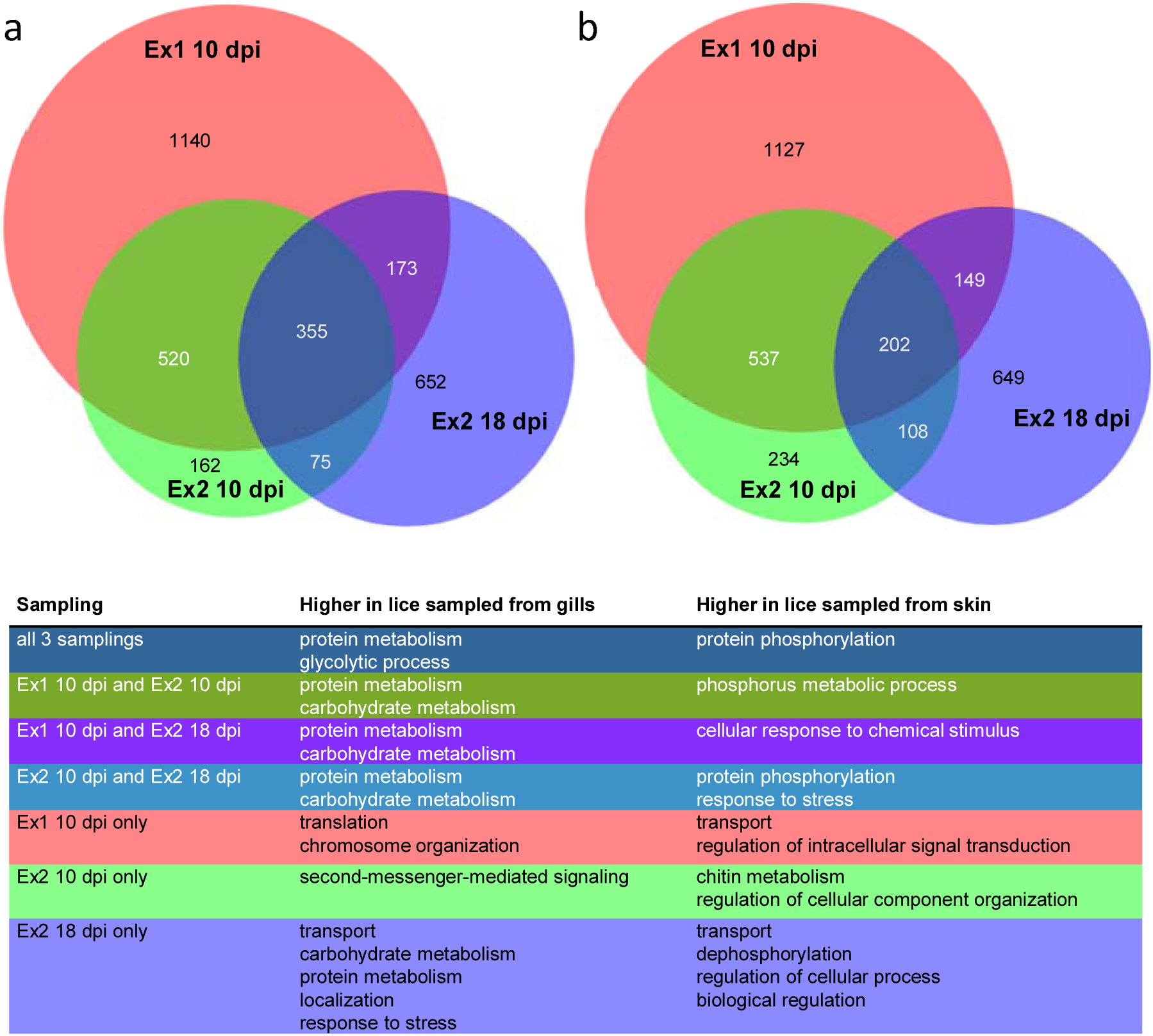
Scaled Venn diagrams showing the number of differentially expressed genes between chalimus larvae sampled from gills versus those sampled from skin, for Experiment 1, sampled at 10 days post infestation (dpi), Experiment 2, sampled at 10 dpi and Experiment 2 sampled at 18 dpi. The numbers of genes elevated in chalimus larvae sampled from gills are shown in (a) and the ones elevated in chalimus larvae sampled from fish skin are shown in (b). Representative GO terms for each group are given in the table in the bottom panel. A full list of enriched GO terms is shown in Supplementary Table S9 which are summarized as Tree maps in supplementary Figure S1.

Among DE genes found at 10 dpi or at 18 dpi we found a high number (51) of genes annotated with Pfam domain PF00040: “Fibronectin type II domain”. Mostly, these are elevated in lice sampled from gills at 10 dpi. However, a smaller number of PF00040 is under the DE genes which are elevated in lice sampled from skin than from gills. Additionally, 85 transcripts with Pfam domain PF00089: “Trypsin” were under the DE genes and are mostly found in the group of lice sampled from gills.

The sizes and overlaps of gene sets that were DE in each experiment separated by expression pattern (elevated in lice sampled from gills or elevated in lice sampled from skin respectively) are depicted in Venn diagrams together with most representative GO terms in Figure 4. All significantly enriched GO terms are listed in Supplementary Table S9. Additional summarized GO annotations belonging to biological process are visualized in a TreeMap (REVIGO) (Supplementary Figure S1). “Peptidase activity” is an enriched GO term in lice sampled from gills across all groups (except the ones exclusively found in Ex2 10 dpi), and in particular “serine type endopeptidase activity”, whereas “serine-type endopeptidase inhibitor activity” is enriched in lice sampled from skin. Notably, “glycolysis” as well as “oxidoreductase activity” are GO terms highly enriched in genes elevated in lice sampled from gills. GO terms containing “phosphorylation” as well as “phosphatase” are found enriched in nearly all groups in genes elevated in lice sampled from skin (also here, the exception is the ones exclusively found in Ex2 10 dpi).

### Equal gene expression changes throughout all three analyses

To determine which genes may be important in relation to the blood meal in general, independent of the stage of the lice at the different time points, we investigated the transcripts which were either significantly elevated in lice sampled from gills or significantly elevated in lice sampled from skin in all three samplings (Ex1 10 dpi, Ex2 10 dpi, Ex2 18 dpi). We found 355 transcripts that were elevated in all three samplings in lice from gills, and 202 from skin. Of the 355 genes elevated in lice sampled from gills, 60% had predicted Pfam domains, and of the 202 elevated in lice sampled from skin, 82% had predicted Pfam domains. A highly prevalent Pfam domain in the DE genes found in all 3 samplings elevated in lice sampled from gills is PF00089: Trypsin. Other more frequent found domains are PF01400: Astacin (Peptidase family M12A), PF02469: Fasciclin domain, PF05649: Peptidase family M13, PF00171: Aldehyde dehydrogenase family, as well as different Zinc finger domains. In the group of DE genes which were elevated in lice sampled from skin, prevalent domains are PF00040: Fibronectin type II domain; PF00069: Protein kinase domain, PF00096: Zinc finger, PF00135: Carboxylesterase family, PF01391: Collagen triple helix repeat. A full list can be found in Supplementary Table S6 and S7. GOslim was used to minimize GO categories. DE genes in lice sampled from gills fall under fewer GOslim categories than DE genes elevated in lice from skin, even though there are more genes in former DE group (Figure 5). Often, genes in the group found elevated in lice sampled from gills fall under the enriched GO group “catalytic activity” (all are shown in Figure 5a). Remarkably strong enriched (factor 30) DE genes elevated in lice from skin, are genes belonging to “extracellular matrix”. However, only five genes are in that group. More than 30 genes were found in GO categories “catalytic activity”, “hydrolase activity”, “binding” and “ion binding” (Figure 5b). All enriched GOslim terms, numbers of genes found in each category, and enrichment factor for the two groups are shown in Figure 5.

**Figure 5.**
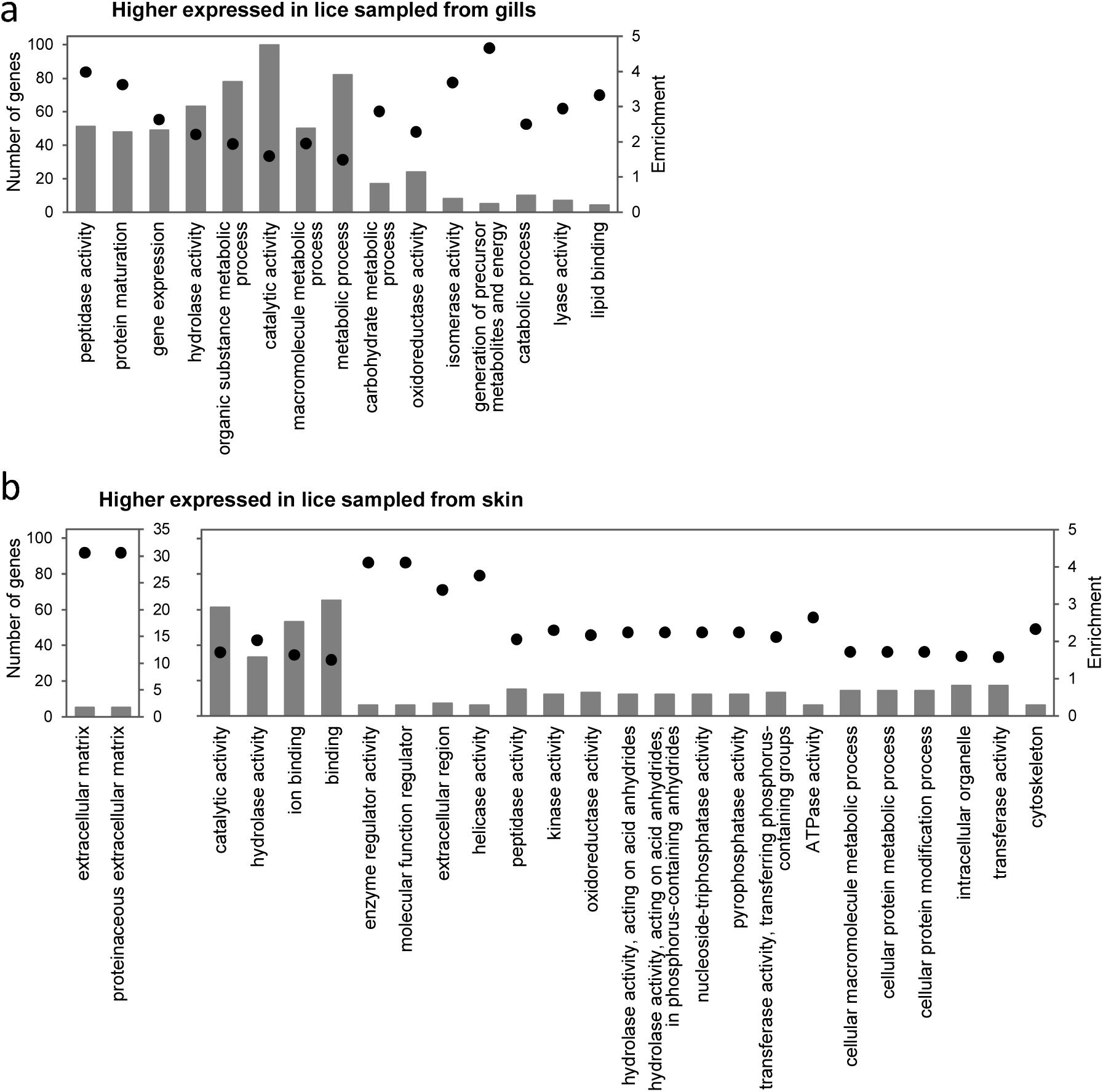
GOslim annotations of genes elevated in lice sampled from gills (a) and genes elevated in lice sampled from skin (b) in all three sampling points. Columns indicate the number of genes in each GOslim category, while black dots show the enrichment of genes in that category in relation to number of genes of the specific category in the whole dataset.

42% of the 355 genes elevated in lice sampled from gills have a fold-change more than two compared to skin. When also taking the p-value into account we find 21%, which are strongly regulated (average fold-change over two and average adjusted p-value ≤ 0.005). In this group, the strongest elevated transcript is most similar to a nematode astacin (EMLSAT00000010457). Among the other strong ones regulated are three transcripts with FNII domains, another transcript with an astacin domain, as well as nine transcripts with trypsin domains, as well as a chemosensory protein (EMLSAT00000005105), yet also many transcripts with no annotation or known protein domains. There are several transcripts with similarity to various proteases that are up-regulated in lice from gills. All genes are listed in Supplementary Table S6. The expression patterns of the ten strongest regulated genes over all samples are shown in Supplementary Figure S2a.

In the group of genes elevated in lice sampled from skin, we find fewer genes highly differentially expressed than in lice sampled from gills. Only 16% have a fold-change of two or higher compared to samples from gills. Most up-regulated in this group is a transcript with no predicted annotation or Pfam domains (EMLSAT00000009920). Among the strongest elevated genes in all three samplings (average fold-change over two and p-value ≤ 0.005) in lice sampled from skin are several transcripts with predicted FNII domains. Moreover, we find several genes with no annotation or known protein domains. All genes are listed in Supplementary Table S7. Expression patterns for the ten strongest regulated ones are shown in Supplementary Figure S2b.

We further looked at the expression profile of these detected DE genes during the course of development, as well as in various tissues (Edvardsen et al., 2014; Eichner et al., 2018; http://licebase.org). We were particularly interested in determining if these transcripts were also elevated in the louse intestine compared with other tissues, or if these transcripts are up- or down-regulated after attachment or after molting to preadult, the expected time point for accessing host blood. Of the 202 genes elevated in lice from the skin, 124 were also investigated in an oligo microarray regarding to expression in different tissues (gut adult female, gut adult male, ovaries, testis, subcuticular tissue and brain) and from the 355 genes elevated in lice sampled from gills 209 were represented in that study (Edvardsen et al., 2014).

Among transcripts elevated in lice sampled from gills, 94 (26%) are higher expressed in the intestine (only 5% of transcripts were lowest in intestine compared with other tissues investigated) (LiceBase). Moreover, 39 of these were very highly expressed in the intestine, compared to other tissues (more than 100 times higher) (LiceBase). 77 of these elevated in intestine were also analyzed in the microarray study investigating different tissues of *L. salmonis* and 52 were also found highest expressed in the intestine there (Edvardsen et al., 2014). 66% of the transcripts are elevated after attachment and 55% are higher expressed in preadult lice than in chalimus II when comparing to the time series data (Eichner et al., 2018). The genes higher expressed in intestine, higher expressed after attachment or higher expressed in pad1 respectively are marked in Supplementary Table S7.

Only 17 (8%) of the transcripts in the DE gene group elevated in lice sampled from skin are higher expressed in the intestine than in other investigated tissues (12% lowest of all tissues investigated) (LiceBase). Only two were much higher (more than 100 times) expressed in intestine than other tissues (LiceBase). However, the ten of these also found on the oligo microarray were not highest expressed in the intestine, except one (EMLSAT00000008355) (Edvardsen et al., 2014). Here two others were found highest expressed compared with the other tissues analyzed. 57% of the transcripts are elevated after attachment. Nearly all of the strongest regulated transcripts are elevated after attachment when comparing with LiceBase data. 53% are higher expressed in preadult lice than chalimus II (Eichner et al., 2018). The genes higher expressed in intestine, higher expressed after attachment or higher expressed in pad1 respectively are marked in Supplementary Table S7.

## DISCUSSION

In this study, we have investigated the biology of blood-feeding in the marine ectoparasitic salmon louse with a special focus on gene expression of immobile lice situated on host gills. We chose immobile lice, because this allowed us to focus on those individuals that had stayed at one location at least since extruding the frontal filament in the late copepodid stage. Being attached to the gills allowed the lice to initiate blood-feeding prior to becoming mobile. Samples from two experiments terminated at 10 days post infestation, and one sample terminated at 18 days post infestation of one the experiments were included in the RNA-seq and subsequent gene expression analyses.

At 10 dpi, the amount of lice is rather evenly distributed between the investigated body parts (32 or 29% on body, 43 or 39% on fins and 24 or 30% on gills in Ex1 and Ex2 respectively). The favored site at 18 dpi is the body with 66 or 61% in Ex1 or Ex2 respectively (15 or 14% on fins and 17 or 23% on gills) (Figure 1). Lice at day 10 are in chalimus I or chalimus II stage and attached by the frontal filament. On the more closely analysed fish at day 18 (Table 2), we found 68 or 66% preadult lice in Ex1 or Ex2 respectively. Of these, 10 or 21% respectively are attached with a filament, while the others are mobile and can freely move on the fish. The finding that lice are differently distributed when being mobile rather than attached, and mainly found on the body of the fish, suggests that the mobile preadult lice choose the general body surface as a preferred feeding site and migrate there from host fins and gills when becoming mobile. The majority (78 or 87% respectively) of the preadult I lice on fins are females. Female lice are known to develop slower than males (Hamre et al., 2019, 2013), and this as well indicates that the lice tend to leave the fins for other host feeding areas when becoming mobile. A preadult I louse that is still attached to its host by its frontal filament has recently molted from the chalimus II stage, and has stayed at that feeding site since attachment. There are no preadult I lice with a visible blood-filled intestine on the fins, whereas we find this on the gills and the body. Interestingly, of the lice still on their filament, only those on gills have apparently fed on blood. Moreover, already in the chalimus I stage, we find lice with blood-filled guts on the gills, but not on any other feeding site. As the preadult I lice on the body with a blood-filled intestine are mobile, these lice have either started with blood-feeding in the mobile preadult I stage or are preadult lice migrated from the gills, meaning that blood-feeding is initiated from the mobile preadult I stage and onwards in the development of the salmon louse occurring under natural conditions.

Development of lice on the gills was delayed, compared to development of lice on body or fins. At 10 dpi, no chalimus II lice were found and a higher proportion of chalimus I lice was of less developed instar age on gills. Comparing only the attached chalimus at 18 dpi (25% of all lice), 51 or 62% in Ex1 or 2 respectively are found on gills. In addition, on gills, there is a higher proportion of male chalimus II lice, which develop faster than females. Developing on host gills caused the salmon lice to develop slower than those developing on other locations. There have been contradicting results about this in the past (Johnson, 1993; Johnson and Albright, 1992), however here we have determined instar ages, and not only the developmental stages, which adds more confidence to our results.

We conclude that during the normal development on the body or the fins, the salmon louse does not start to feed on blood until reaching the mobile preadult I stage. By that reasoning, we wanted to compare gene expression of chalimus larvae located on the vascular gills with access to blood with that of chalimus larvae of equal development from the rest of the body. The salmon louse has approximately 13,000 protein encoding genes (http://metazoa.ensembl.org/Lepeophtheirus_salmonis), and we find in our RNA-seq analyses that over 5800 genes had an altered expression in at least one of our samplings. As expected, we found a higher amount of overlapping DE genes in the two samplings at 10 dpi. These are chalimus I larvae, which are soon molting to chalimus II, while lice sampled at 18 dpi are chalimus II larvae, shortly prior to molting to preadult I lice. As such, all lice are sampled at a similar instar age. However, phenotype and lifestyle differ in preadult lice and one can expect expression of genes in preparation for this stage in the lice sampled at 18 dpi. The high amount of DE genes exclusively found in Ex1 10 dpi could be caused by batch differences between Ex1 10 dpi and Ex2 10 dpi, or could be as a result of more powerful statistics due to a higher amount of parallel samples (8 versus 6 biological parallels of each group in Ex1 10 dpi and Ex2 10 dpi, respectively). However, we know also that minor differences in development have a high impact on gene expression (Eichner et al., 2018), and individual differences occurring within groups, with possible consequences between groups, could bias the results.

To investigate gene expression caused by nutrition differences, we mainly concentrated on the DE genes found in all 3 samplings. Transcripts over-expressed in lice sampled from gills could be important for hematophagy. However, many (70 of 74) of the strongest DE genes in this group are not highest expressed in the intestine, but rather in other tissues, suggesting these contribute to other functions in the louse that may be modified by hematophagy. Genes elevated in lice from gills show a more homogenous GO annotation (fewer GOslim categories) than the ones elevated in lice from skin, suggesting that several DE genes are involved in the same processes. There are also more genes with a greater fold change within the group of DE genes elevated in lice sampled from gills (42% over 2-fold change, whereas only 16% in lice sampled from skin), pointing towards a high demand of these gene products when feeding on blood. However, as GO terms can be unspecific or general, the following discussion deals with selected groups of transcripts.

### Iron and heme

Among the regulated transcripts, the iron storage units of ferritin (*LsFer1* (RACE sequence) and *LsFer2*: EMLSAT00000006305) are both elevated in chalimus larvae sampled from gills compared with other settlement sites (Supplementary Figure S3 a, b). We have previously established that these genes are important for the adult female salmon louse blood-feeding and reproductive success, as the parasite had a clear gut and failed to produce viable eggs upon silencing the two genes (Heggland et al., 2019b). Blood contains several iron-proteins, and when initiating blood-feeding, the salmon louse needs to obtain a way of storing and detoxifying iron absorbed from the blood. Up-regulating ferritin when ingesting a blood meal is therefore an important defense mechanism for a blood-feeding parasite. The putative heme scavenger receptor, *LsHSCARB* (EMLSAT00000005382), is elevated in lice on gills at 18 dpi compared to lice on skin (Supplementary Figure S3d). We recently found that upon silencing *LsHSCARB* by RNA interference, adult female lice had absorbed less heme and produced fewer viable eggs and less offspring (Heggland et al., 2019a). Lacking early (10 dpi) transcriptional elevation of *LsHSCARB* could indicate alternative mechanisms of absorption during the earlier developmental stages, or the existence of a post-transcriptional mode of regulating the LsHSCARB protein. Alternatively, the lack of early regulation might serve to maintain homeostasis of heme levels when feeding on the vascular gills.

### Detoxification

A Glutathione S-transferase (GST) (PF02798) transcript (EMLSAT00000009830) was elevated in lice on gills in all samplings. GSTs are major detoxification enzymes. A GST in the hard tick *Ixodes ricinus* (IrGST1) (GenBank ID: MF984398) was also found to be elevated in the midgut of blood-fed ticks compared with serum-fed ticks (Perner et al., 2016). Further characterization of IrGST1 showed that it was heme-inducible and the recombinant protein was able to bind heme *in vitro* (Perner et al., 2018). The authors speculated that IrGST1 is important for detoxifying excess heme to avoid cytotoxicity in the tick (Perner et al., 2018). Recombinant GSTX2 (GenBank ID: AAK64286.1) of *A. aegypti* also binds heme (Lumjuan et al., 2007), and was elevated in a heme-incubated *A. aegypti* Aag2 cell line (Bottino-Rojas et al., 2015). Of the six different predicted salmon louse proteins with the GST domain (PF02798), EMLSAP00000009830 is the most similar to both IrGST1 and *A. aegypti* GSTX2. The connection of GST and blood-feeding in the salmon louse is an interesting topic for future studies, as we per today do not know what mechanisms the salmon louse depends on to detoxify heme.

### Digestion

Food protein hydrolysis is a fundamental step of digestion, and is mediated by peptidases that enzymatically cleave peptide bonds. Blood is highly enriched in protein, and one of the most abundant ones is the gas transporter hemoglobin. Investigating changes in the salmon louse transcriptome upon initiating blood-feeding could thus give clues as to which enzymes are essential for breakdown of blood components. Trypsin is a digestive enzyme belonging to the S1A subfamily of serine endopeptidases, and five main trypsin-encoding transcripts in the salmon louse intestine have previously been characterized (Johnson et al., 2002; Kvamme et al., 2004). Trypsins and other proteins involved in protein degradation were found elevated e.g. in blood-fed mosquito *A. aegypti* (Bonizzoni et al., 2011). Twenty-eight transcripts with trypsin as the only predicted protein domain (PFAM: PF00089) (29 in total with trypsin + other domains) were found to be elevated in lice on host gills at day 10 (Ex1 and 2) and 18 dpi. Of these, 11 are predicted to be highest expressed in the intestine compared with other tissues investigated in the salmon louse (LiceBase; Edvardsen et al., 2014; Supplementary Table S6). A heat map showing the expression patterns for all transcripts with trypsin domains found DE in all 3 samplings in data taken from LiceBase and from the time-series study are shown in a hierarchical cluster in Figure 6. *LsTryp1* (GenBank ID: AY294257, best blast hit: EMLSAT00000004828) is elevated in all three samplings in lice on gills. One transcript with a trypsin domain only (EMLSAT00000004988) is found to be elevated in lice on host skin at both 10 dpi (Ex1 and Ex2) and at 18 dpi. RNA-seq data in LiceBase as well as microarray data from Edvardsen et al (2014) however show that this transcript has a low expression in the louse intestine and is rather expressed in antenna and feet (LiceBase) or subcuticular tissue and brain (microarray). It might therefore be of importance for other purposes than blood meal digestion.

**Figure 6.**
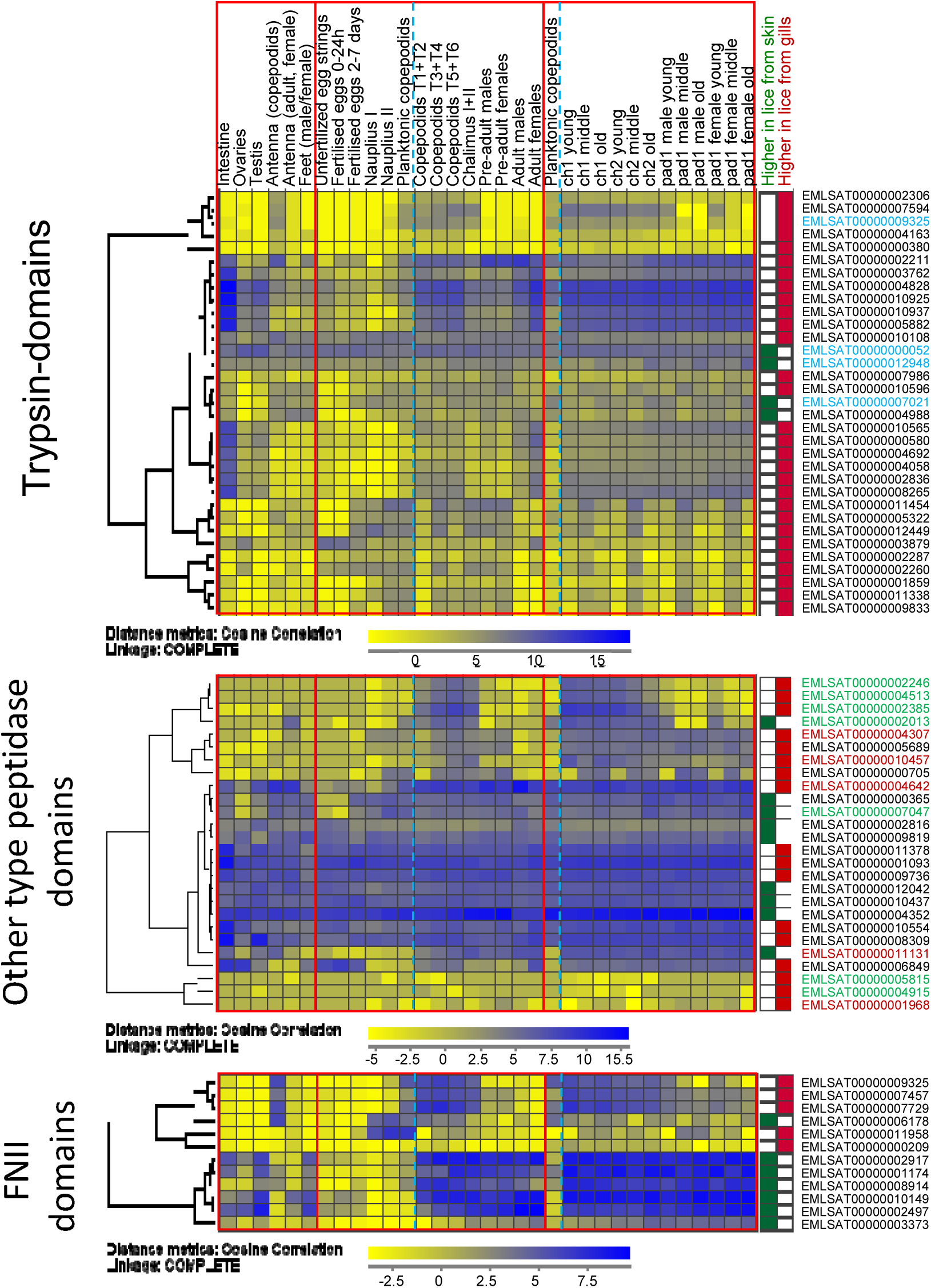
Hierarchical clustering of the expression profiles of genes found DE in all three sampling points with trypsin, other peptidase and FNII domains after expression profiles from LiceBase (various tissues and stages) and from the time series study (average values of biological parallels) by Eichner et al. (2018). A blue stippled line is separating planktonic and parasitic stages. Stable IDs with font in blue = domains other than trypsin predicted as well, font in green = predicted M13 peptidases, font in red = predicted Astacin peptidases.

Peptidases other than trypsins are also regulated in lice on host gills. There are 17 transcripts with Pfam domains “peptidase” other than trypsins elevated in lice on gills in all samplings. Among these are four transcripts with Astacin-domains (Peptidase family M12A) and five are M13 peptidases (Figure 6). Both groups are metallopeptidases and are enriched in arthropods. Astacin-like metallopeptidases are implicated in digestive processes, but are also reported to have anticoagulative effects, as they are found to have fibrinogenolytic activity in spider venoms (Trevisan-Silva et al., 2010). M13 metallopeptidases are widely distributed in animals, and e.g. make up the major group of the hematophagous tick degradome (Mulenga and Erikson, 2011). Furthermore, we also find many of the same types of peptidases elevated rather in lice from skin (one with Astacin domain, two with Peptidase family M13 domain). This could indicate different modes of digesting a blood meal versus digesting components of ingested salmon skin. Further investigation into the elevated trypsins and other peptidases expressed in the salmon louse gut should be conducted.

### Putative anti-coagulation

Blood coagulation is a key mechanism in maintaining homeostasis in vertebrates if a blood vessel were to rupture. A parasite feeding on vertebrate blood would therefore require mechanisms in order to counteract this to maintain its feeding habit. Anti-coagulation factors targeting host proteins could thus be vital for the successful blood-feeding in the parasitizing arthropod. A thrombin (coagulation factor) inhibitor, hemalin, was found to be important to avoid clotting of the blood meal in the bush tick *Haemaphysalis longicornis* (Liao et al., 2009). A salmon louse transcript (EMLSAT00000003009) encoding two Kunitz/Bovine pancreatic trypsin inhibitor domains (PF00014), as also found in the tick hemalin, was elevated here in all three samplings in lice on gills. However, four other transcripts with the same domain were elevated in lice from skin in all samplings (EMLSAT00000000152, EMLSAT00000007907, EMLSAT00000008877 and EMLSAT00000009255).

We also find serine protease inhibitors (serpins, PF00079) regulated. From the 15 predicted serpin transcripts in the louse, four were DE in all three samplings. Two were elevated in lice on gills (EMLSAT00000010931 and EMLSAT00000001743), one elevated in lice on skin (EMLSAT00000011353), while the last (EMLSAT00000005224) was expressed lower at 10 dpi, but elevated at 18 dpi in the lice sampled from gills. One transcript (EMLSAT00000000552) was elevated in lice on gills at 10 dpi (Ex1 and Ex2). Anti-coagulation factors could be targets for pest control as they are likely secreted and in contact with the host, and thus probably vital for the host-parasite interaction.

### Fibronectin type II

The approximately 60 amino acid long fibronectin type II (FNII) domain (PF00040) is a protein domain found within the glycoprotein fibronectin. It contains four conserved cysteine residues that form disulfide bridges. These residues are important for e.g. fibronectin’s collagen binding properties (Guidry et al., 1990). The FNII domain is also found within the vertebrate blood coagulation protein Factor XII (McMullen and Fujikawa, 1985). The FNII domain is the most expanded protein domain of the salmon louse with over 200 copies within over 80 genes identified so far (http://metazoa.ensembl.org/Lepeophtheirus_salmonis). Some of these genes (*LsFNII1, 2* and *3*) have been characterized, and are expressed in tegumental type 1 (teg1) glands of the salmon louse (Harasimczuk et al., 2018; Øvergård et al., 2016). Teg1 glands are exocrine and their secretory ducts are extending to the dorsal and ventral side of the salmon louse (Øvergård et al., 2016). The functions of FNII-containing proteins have not been determined in the salmon louse, however it has been suggested that proteins with the domain may be of importance for lubricating the integument and functioning as an anti-fouling agent, or as part of the salmon louse fuzzy coat (acid mucopolysaccharide layer (Bron et al., 2000)) (Harasimczuk et al., 2018). Genes with FNII domains expressed in teg1 glands have also been suggested to be of importance for host immune modulation by the parasite (Øvergård et al., 2016).

We find that several FNII-containing genes are significantly regulated in chalimus larvae in our dataset. Five transcripts with predicted FNII domains are elevated in all three samplings in lice on gills. Of these, all but one (EMLSAT00000011958) are predicted to be up-regulated after louse attachment (http://licebase.org; Supplementary Table S6) (Figure 6). Seven transcripts with FNII are rather found elevated in lice on skin in all three experiments. Here as well, all but one transcript (EMLSAT00000006178) are up-regulated after louse attachment (http://licebase.org; Supplementary Table S7). The FNII encoding genes characterized by Øvergård et al. (2016) and Harasimczuk et al. (2018) are not among the transcripts regulated in all samplings here (*LsFNII1*: EMLSAT00000012082, *LsFNII2*: EMLSAT00000007294, *LsFNII3*: EMLSAT00000009744), however LsFNII1 was elevated in both experiments at 10 dpi, and LsFNII3 at 18 dpi as well as in Ex1 10 dpi. The transcripts elevated in lice on gills should be further characterized, in order to elucidate a possible role of FNII in blood-feeding. Given the earlier reports that FNII domains in vertebrates may be important for blood clotting, one hypothesis is that proteins with FNII domains only could have an anti-coagulant effect.

## CONCLUSIONS

Blood is a major dietary component for the ectoparasitic salmon louse, which the parasite has access to when attached to a salmonid host. We find that the salmon louse initiates blood-feeding during the mobile preadult I stage. However, if the parasite is attached to host gills, it may start feeding on blood already at the chalimus I stage or even earlier. Blood can be found even in copepodids sampled from gills (personal observation, Supplementary Figure S4). The premature onset of blood-feeding caused lice on gills to develop at a slower pace than lice that were attached to host fins and general body surfaces. Chalimus lice of equivalent age on gills versus other attachment sites were therefore analyzed for gene expression comparisons. Several genes are elevated in lice attached to the gills, and among these, we find e.g. genes of importance for the absorption, storage and/or transportation of the pro-oxidative molecules iron and heme, digestive and detoxification enzymes, genes that could be important for anti-clotting of host blood, and several genes with FNII domains. The results of this study raise a number of new gene targets to investigate further in order to elucidate the blood-feeding habit of the infamous salmon louse.

## ACKNOWLEDGEMENTS

This research has been funded by The Research Council of Norway, SFI-Sea Lice Research Centre, grant number 203513/O30 and 226266. Further, this work was funded by the ELIXIR2 (270068) infrastructure grant from the Research Council of Norway to MD. The sequencing service was provided by the Norwegian Sequencing Centre (www.sequencing.uio.no); a national technology platform hosted by the University of Oslo and supported by the “Functional Genomics” and “Infrastructure” programs of the Research Council of Norway and the South-Eastern Regional Health Authorities. We would like to thank Lars Are Hamre and Per Gunnar Espedal for help in the wet-labs and Madeleine Carruthers for helpful comments on the manuscript.

## AVAILABILITY OF DATA

The datasets supporting the conclusions of this article are included within the article and its additional files. Raw RNA-sequencing data files have been deposited to NCBI BioProject under the accession number PRJNA577842. A preprint of this manuscript and its supplementary files have been made publicly available on bioRxiv: https://doi.org/10.1101/815316.

---

**Supplementary Fig S1.**
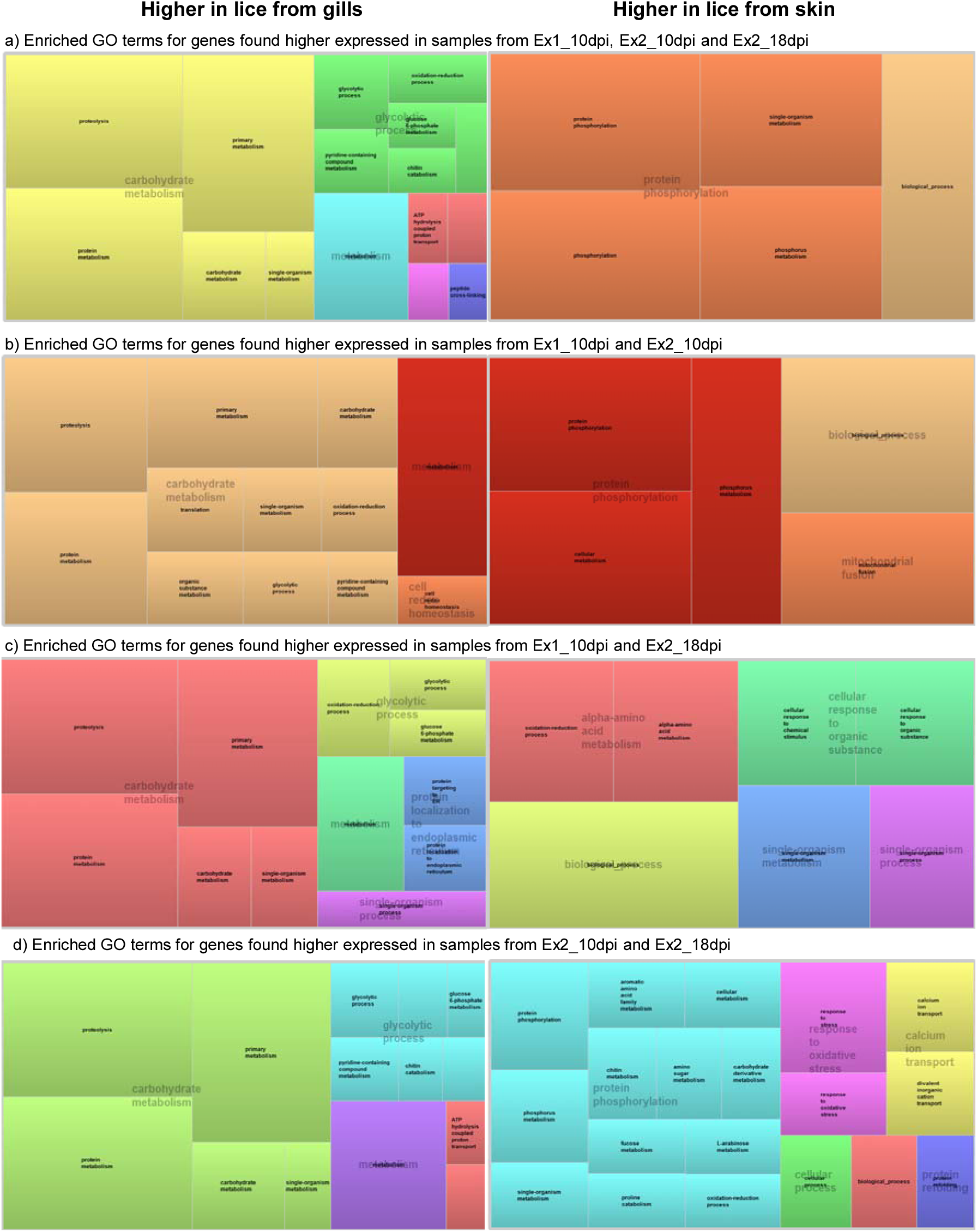
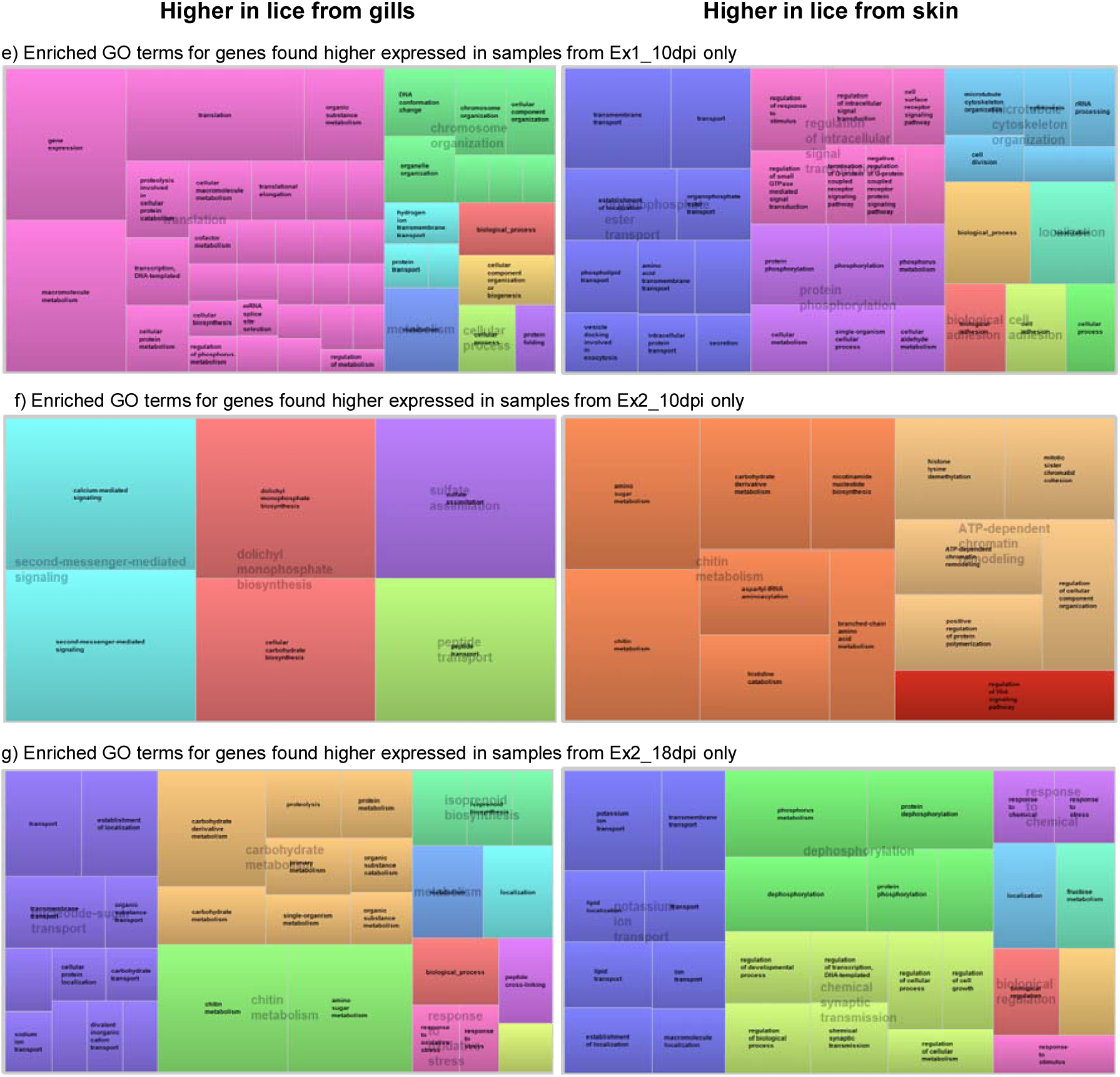
Tree maps (Revigo) of GO annotation belonging to biological process for a) enriched GO terms for genes found higher expressed in samples from all three samplings; b) of the genes DE in both samplings at 10 dpi; and in 2 samplings sampled at different time point (c and d). In e-g are enriched GO annotations shown for genes differentially expressed in one of the three samplings only.

**Supplementary Fig S2.**
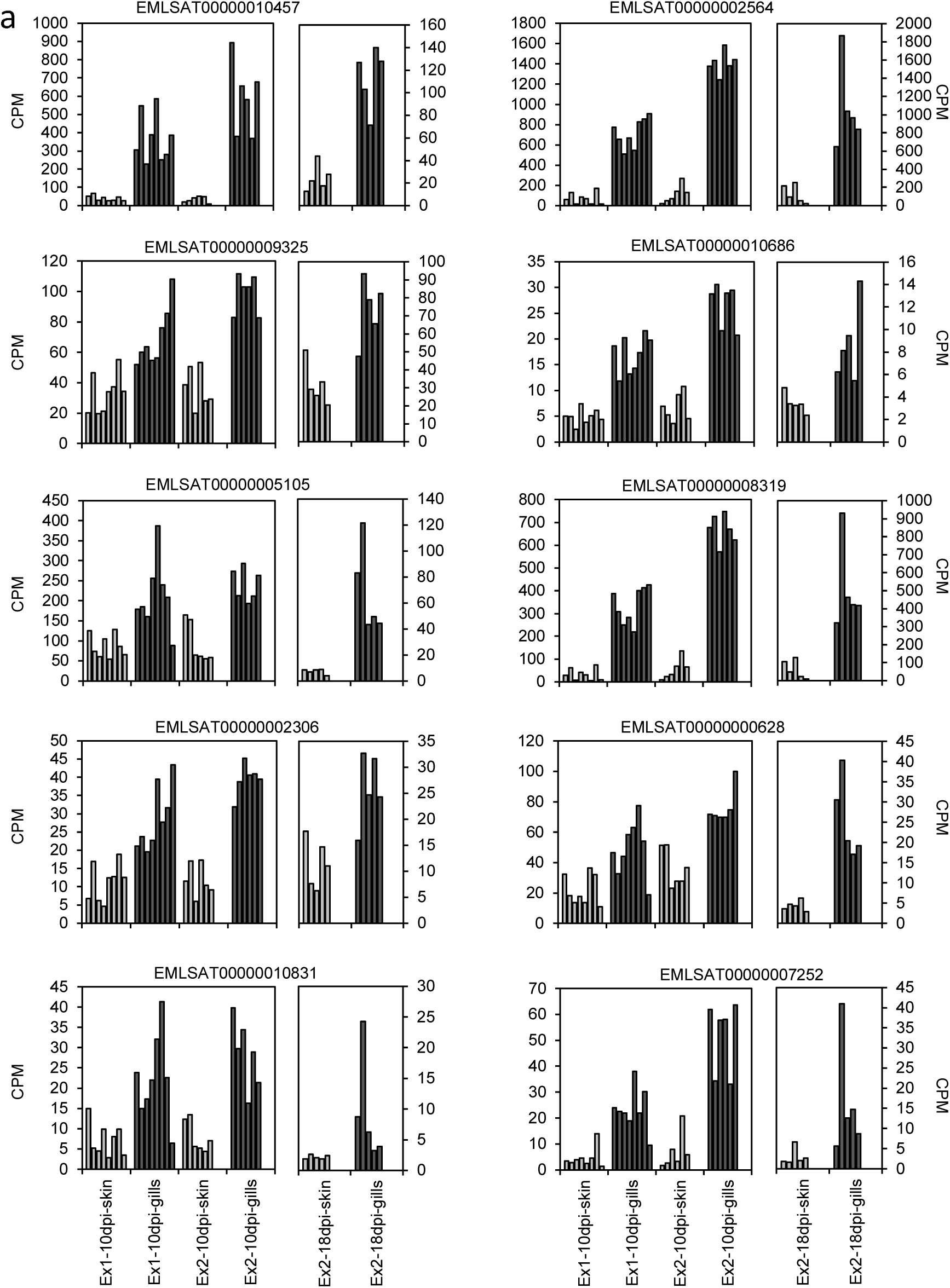
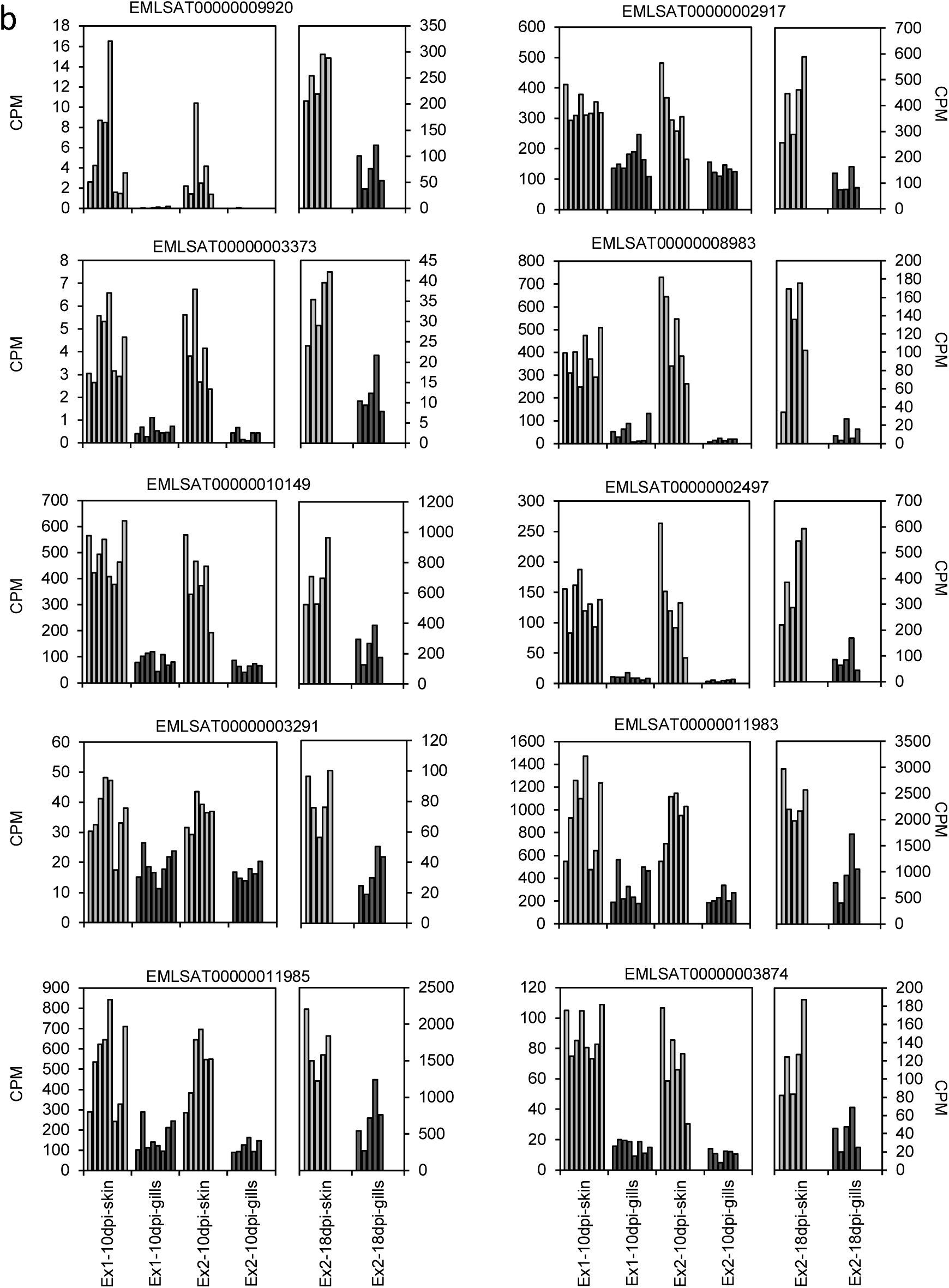
Expression profiles of the ten strongest genes upregulated in lice sampled from gills (a) and in lice sampled from skin (b) over all samples.

**Supplementary Figure S3.**
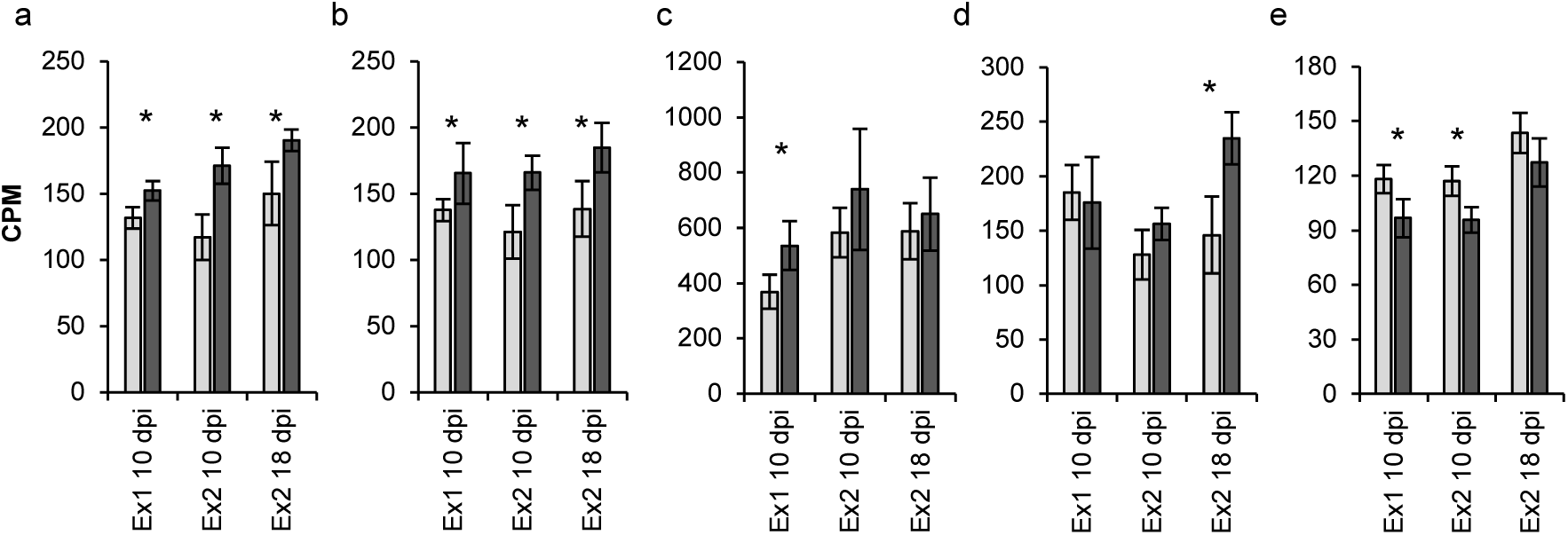
Expression of a) Ferritin 1, b) Ferritin 2, c) Ferritin 4, d) LsHSCARB, e) Lipid transfer protein (EMLSAT00000001530)(Khan et al., 2017) in the samples from lice sampled from skin (light grey) and gills (dark grey). Shown are average values with standard deviation. * = Significant different (T-test) between lice samples from different location in the respective sampling point.

**Supplementary Fig S4.**
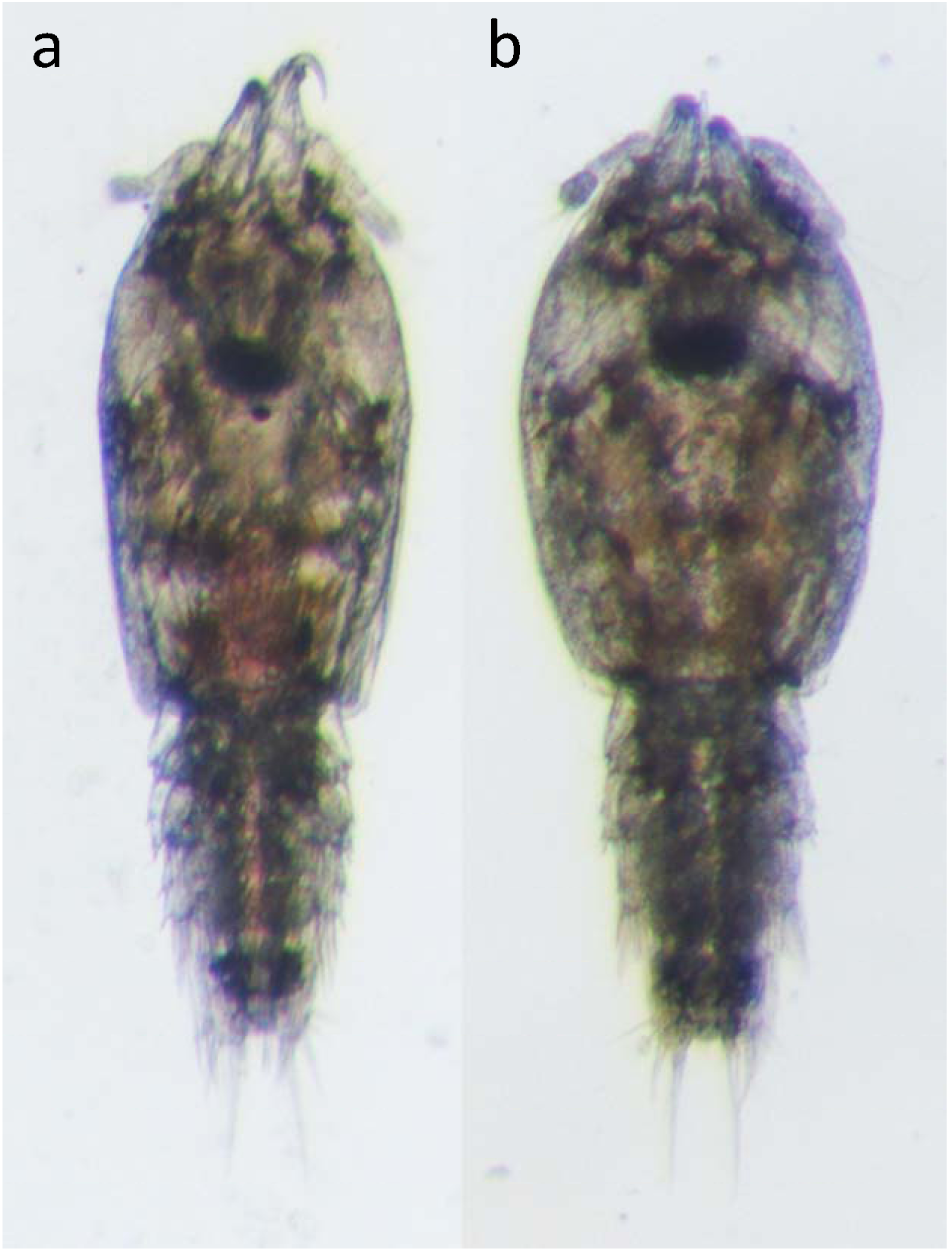
Copepodite with blood in intestine sampled from gills (a) and without blood (b) at 3 days post infestation (samples not included in this study).

